# Hypothetical lytic transglycosylase SleB is important for cell fitness in *Zymomonas mobilis*

**DOI:** 10.64898/2026.07.21.739787

**Authors:** Katsuya Fuchino, Richard Daniel, Christodoulos Astraios, Waldemar Vollmer

## Abstract

Bacterial peptidoglycan (PG) undergoes a variety of chemical modifications. O-acetylation at the C6 hydroxyl group of *N*-acetylmuramic acid is a widespread PG-modification found across diverse bacterial phyla. It contributes to virulence in pathogenic bacteria because the O-acetyl group reduces the activity of the PG-degrading host defense enzyme, lysozyme. Beyond its role in host defense evasion, recent studies suggest that PG O-acetylation also regulates the activity of endogenous lytic transglycosylase (LT) autolysins.

The ethanologenic alpha-proteobacterium *Zymomonas mobilis* O-acetylates its PG, which is associated with tolerance to environmental stresses, including salt. To better understand how PG O-acetylation contributes to stress tolerance, we investigated the predicted lytic transglycosylase SleB. Intriguingly, the *sleB* gene is located adjacent to the *pat* operon, which encodes the proteins responsible for PG O-acetylation. We showed that loss of SleB caused impaired growth and morphology, and a significant reduction of crosslinks in the PG of *Z. mobilis.* Furthermore, the *sleB* mutant was sensitive to environmental stress resembling the sensitivity of the *patA* mutant. Collectively, our findings unravelled an important role of SleB in PG remodelling and stress resilience.

## Introduction

Bacteria surround their cytoplasmic membrane by peptidoglycan (PG), which forms a rigid and elastic macromolecular sacculus made of glycan chains of *N*-acetylmuramic acid (Mur*N*Ac) and *N*-acetylglucosamine (Glc*N*Ac) repeats and peptides, which can be cross-linked (1). While this mesh-network structure provides mechanical strength and protects cells from disruption due to its turgor, the cell needs to constantly remodel PG for growth and cell division.

PG remodelling is mediated by a diverse group of enzymes collectively referred to as PG hydrolases, which cleave specific bonds within the PG (2). These activities are essential for making space in the PG for the insertion of newly synthesised PG into the existing layer, regulation of PG growth and PG remodelling for insertion of cell envelope complexes such as flagella.

Lytic transglycosylases cleave glycan strands in PG (3) to generate muropeptides with a 1,6-anhydro-MurNAc residue, which is typical for PG turnover products (4) and glycan chain termini (5). LTs also remove excessive PG that accumulates in the periplasm (6) or in the presence of beta-lactam antibiotics (7). LT activity is affected by PG O-acetylation (8), the chemical modification that adds an acetate to the C6 hydroxyl group of MurNAc. This modification inhibits exogenous lysozyme and contributes to the resistance to this antibacterial, PG-degrading enzyme in Gram-positive pathogens such as *Staphylococcus aureus* (9) or *Streptococcus pneumoniae* (10). In Gram-negative bacteria, the PG is protected from exogenous lysozyme by the outer membrane (11), yet some species O-acetylate and de-O-acetylate their PG to control their endogenous LT enzymes (12). In Gram-negative bacteria, O-acetylation is mediated by O-acetyltransferase PatA and PatB, while the esterase Ape removes the acetate group from the PG (12, 13).

The alpha-proteobacterium *Zymomonas mobilis* is well known for its ethanologenic physiology (14) and thus a promising species for industrial applications (15, 16).

However, its sensitivity to environmental stresses, including salt, remains a bottleneck for industrial applications. We recently showed that engineering PG remodelling can improve resilience to salt stress, underscoring the importance of studying the cell envelope or this organism (17). We also discovered that *Z. mobilis* O-acetylates its PG and that this modification is important for growth under salt stress (17). These findings suggest that the robustness to stress of *Z. mobilis* may be improved by manipulating its PG structure.

Here, we investigated the role of SleB, whose gene locates next to the *pat* operon. The obtained data shows that SleB significantly contributes to cell growth, morphology and stress resilience in *Z. mobilis* cells.

## Methods

### Bacterial strains, plasmids, and growth conditions

Bacterial strains and plasmids used in this study are shown in Table S1. *Z. mobilis* was grown in RM complex medium containing glucose (20 g/L), yeast extract (5 g/L), NH_4_SO_4_ (1 g/L), KH_2_PO_4_ (1 g/L), and MgSO_4_ (0.5 g/L). The complex medium was briefly sparged with nitrogen gas to make it micro-aerobic. Twelve-mL cultures of *Z. mobilis* were incubated at 30°C in a capped 15 ml Falcon tube without shaking. For obtaining the growth profiles, the optical density was measured in a 96- well plate by SPECTROstar Nano (BMG labtech).

### Construction of *Z. mobilis* strains

We employed a published method for genetic manipulations (18). To construct plasmids for genetic engineering, the upstream and downstream regions of the gene of interest were amplified by polymerase chain reaction (PCR) using Q5 DNA polymerase (New England Biolabs). The obtained fragments were inserted into the plasmid pPK15534 by Gibson Assembly (New England Biolabs). The sequence of constructed plasmids was verified by Sanger sequencing. The plasmids were introduced into *Z. mobilis* strain ZM6 via conjugation with *Escherichia coli* strain WM6026 as described (19). *Z. mobilis* cells with the integrated plasmid in their chromosomal DNA were selected by 90 μg/mL chloramphenicol in RM medium agar. The integration of the plasmid into the chromosomal DNA was confirmed by colony PCR. For excision of the plasmid, the *Z. mobilis* cells were grown overnight in liquid RM medium without selection, and subsequently the loss of plasmid was achieved by counterselection with the *Bacillus subtilis* SacB that had been optimised for *Z. mobilis* (18).

For cloning of *pdc*-*sleB* into the plasmid pBBR, the *sleB* gene and promoter region of *pdc* were amplified by Q5 DNA polymerase. These two fragments were annealed using overlap PCR by Q5 DNA polymerase. The annealed product and the plasmid pBBR were cut by the restriction enzyme XbaI and XhoI, and purified using a PCR purification kit (Qiagen). These two products were ligated by T4 ligase (New England Biolabs) and transferred into *Z. mobilis* strains via conjugation. Ex-conjugants were selected on RM medium containing kanamycin (220 μg/mL).

The primers used for constructing and verifying the strains are listed in Table S2. The genomic DNA of Δ*sleB* and Δ*mltA* was extracted using a genomic DNA kit (Sigma) and sequenced by whole genome sequencing (Illumina MiSeq, Illumina). The genotype, including heterogeneity variants ratio of specific mutation, was analysed using CLC Genomics Workbench version 25.0.5. (Qiagen).

### Microscopy and image analysis

A culture with growing *Z. mobilis* (0.5 µl) was spotted onto a 1% agarose-pad made of RM growth medium and covered by a cover-glass. Phase contrast and fluorescence images of *Z. mobilis* cells were taken using a Nikon Eclipse Ti microscope (Nikon) equipped with a Plan Apochromat 100x objective with an NA of 1.40 and Ph3 Phase plate (Nikon) and a Prime sCMOS camera (Teledyne Photometrics). Images were acquired using Nikon NIS elements AR software. Image analysis was performed using MicrobeJ (20). Unpaired T-test was applied to assess statistical significance.

### Muropeptide analysis

A *Z. mobilis* culture (400 mL) was anaerobically grown to an optical density of 0.6 –0.8. The cells were harvested by centrifugation, resuspended in ice-cold Phosphate-Buffered Saline (PBS) and subsequently boiled in 4% sodium dodecyl sulfate (SDS). Purification of peptidoglycan and high-performance liquid chromatography (HPLC) was performed as described (17).

## Results

### *Z. mobilis* Δ*patA* is sensitive to SDS and acetic acid stress

We have previously shown that the PG of *Z. mobilis* is O-acetylated and that the deletion of the *patA* gene encoding PG O-acetyl transferase abolished this modification (17). Interestingly, the *patA* mutant (Δ*patA*) was shown to be sensitive to salt stress (225 mM NaCl) without exhibiting an apparent growth defect under regular growth conditions (17). The mechanism underlying the salt sensitivity is not yet known. As the salt sensitivity is likely an indirect effect of the absence of PG O-acetylation, we hypothesized that Δ*patA* might also be sensitive to other stresses.

We first examined sensitivity to sodium dodecyl sulfate (SDS) to assess whether the loss of PG O-acetylation compromises membrane integrity. Wild-type ZM6 cells grew slightly slower when SDS was present. Interestingly, we found that Δ*patA* was sensitive to SDS, growing slower and reaching lower final biomass than the wild-type (Fig. 1B), suggesting that the cell envelope structure and integrity is compromised in the mutant.

**Fig. 1.**
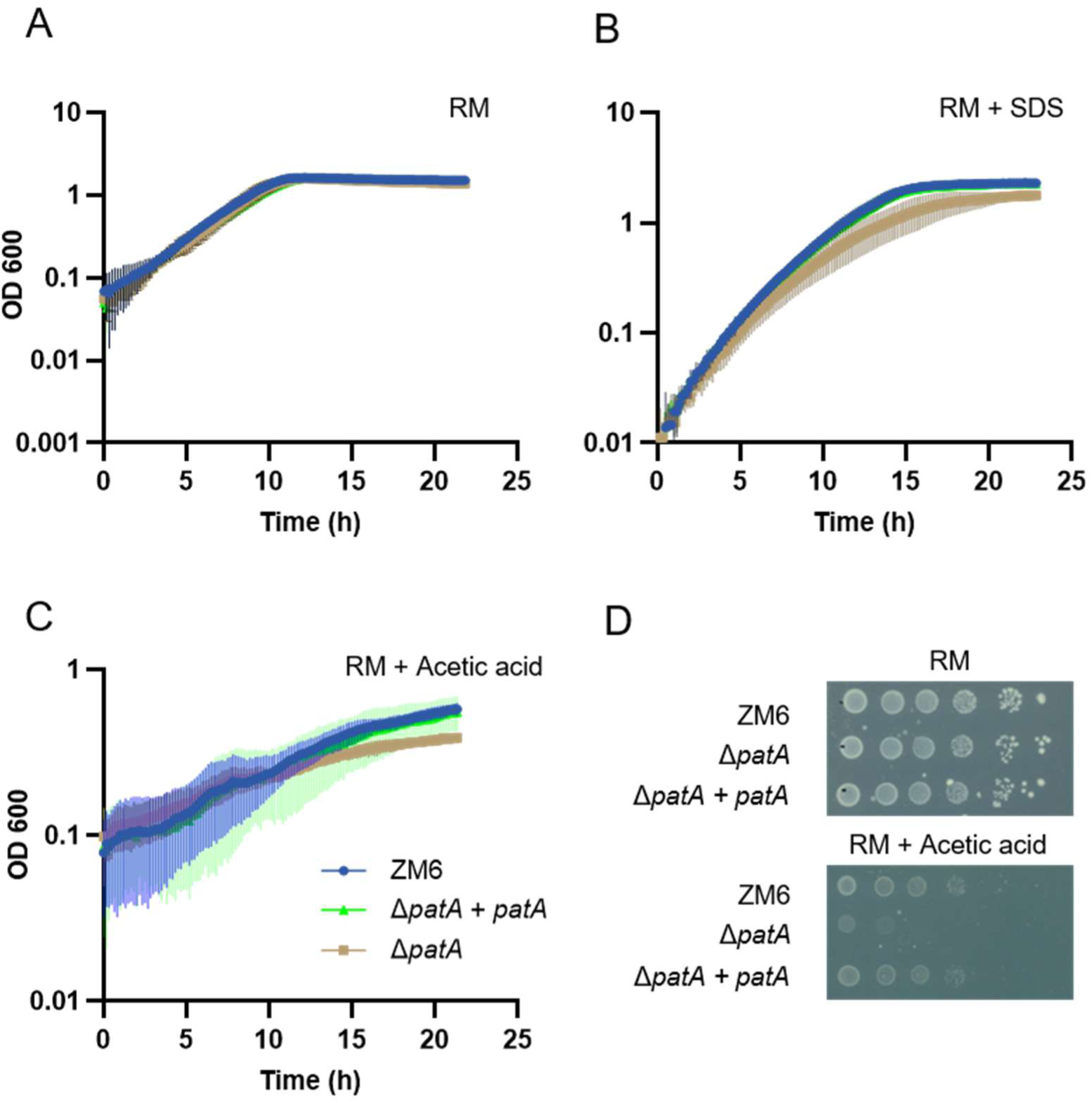
Δ*patA* is sensitive to SDS and acetic acid stress. Growth of *Z. mobilis* strains ZM6 (wild-type), Δ*patA*, Δ*patA + patA* (complementation strain) under regular conditions **(A)**, or in the presence of SDS (0.0025%) **(B)** or acetic acid **(C)**, measured in a 96-well plate reader. **(D)** The spot assay was performed to confirm the acetic sensitivity of Δ*patA.* The image was taken after 96 h. Biological replicates, N = 3. The error bars represent the standard deviation.

We next examined the effect of environmental acetic acid on the Δ*patA* mutant. Notably, acetic acid stress is also one of the most notorious stresses that hindered the use of *Z. mobilis* in biofuel production. Supplementing the wild-type strain with acetic acid (3.5 g/L, 58 mM) impaired growth as previously shown (21). Interestingly, Δ*patA* was even more sensitive to acetic acid than the wild-type (Fig. 1CD). These results, together with the previously observed salt sensitivity, indicate that in the *Z. mobilis* PG O-acetylation contribute to cell envelope stress resilience.

### Δ*sleB* has growth and morphology defects

To understand the phenotype of the PG O-acetylation deficient Δ*patA* strain, we considered that the observed phenotype might arise from unregulated PG remodelling. PG O-acetylation is known to inhibit endogenous LTs and thus, the absence of O-acetylation could result in elevated LT activity. Interestingly, one of the adjacent genes of the *pat* operon is annotated as *sleB* and encodes a soluble LT (Fig. 2A). This genetic link indicates that the function of both gene products might be linked. We generated the mutant lacking the putative LT gene (Δ*sleB*) using an optimised homologous recombination method we previously developed (18, 19). As *Z. mobilis* is a polyploid species (22–24), the mutation could result in a heterozygous genotype. Thus, we verified the mutation by whole genome sequencing and the deletion on all chromosome copies (18).

**Fig. 2.**
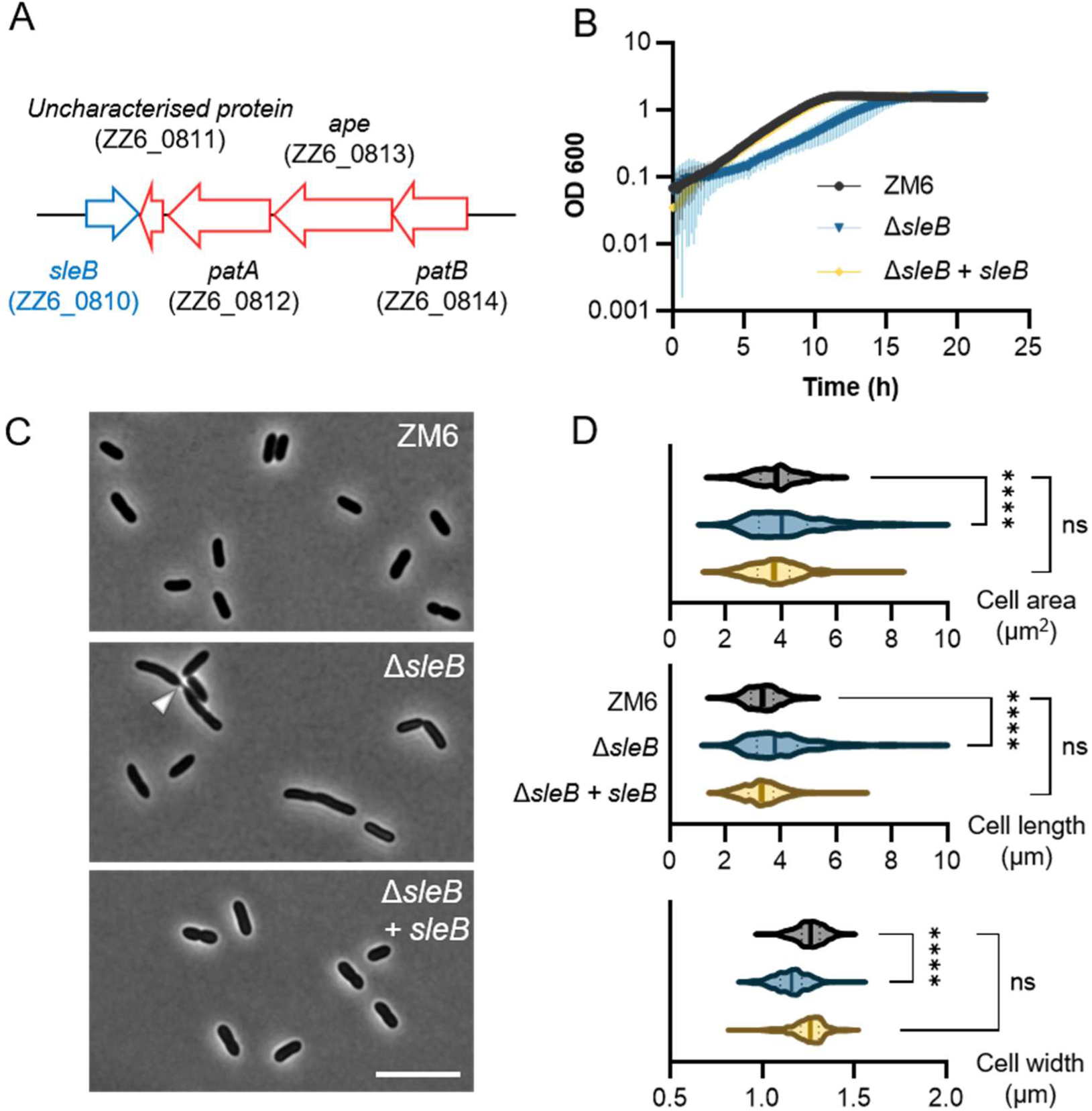
*Z. mobilis* Δ*sleB* cells exhibit impaired growth and morphology defects under regular growth conditions. **(A)** A schematic representation of the neighbourhood of the *pat* operon with the *sleB* gene. **(B)** Growth of *Z. mobilis* strains ZM6 (wild-type), Δ*sleB*, Δ*sleB* + *sleB* (complementation strain) under regular growth conditions. Biological replicates N = 3. The error bars represent the standard deviation. **(C)** Phase contrast images of growing ZM6, Δ*sleB*, Δ*sleB* + *sleB* under regular growth conditions. An arrow head indicates incomplete cell separation in Δ*sleB* cells, showing a thin thread-like structure connecting the two daughter cells. Scale bar, 10 μm. **(D)** Box-and-whisker plot showing the area, length and width of actively *Z. mobilis* cells (OD 0.5 - 1.3) of the three strains under regular growth condition, measured using MicrobeJ (20). Biological replicates N = 3. Sample size, n = 459 cells (ZM6), n = 447 cells (Δ*sleB*), n = 435 cells (Δ*sleB + sleB*).

We next compared growth and cell morphology of the Δ*sleB* deletion strain and the parental strain ZM6 under regular growth conditions. The Δ*sleB* cells grew significantly slower than the wild-type, although the Δ*sleB* culture reached a final optical density comparable to that of the wild type (Fig. 2B). The wild-type cells were rod-shape during active growth, but the Δ*sleB* cells were thinner and longer and occasionally formed cell chains (Fig. 2CD). To exclude potential polar effects, we complemented the mutant by inserting *sleB* with its upstream region into a different locus of the chromosome. The complementation restored the WT phenotype, verifying that the observed growth defects were due to the *sleB* mutation (Fig. 2BCD).

Many bacteria have several, functionally redundant LTs (6) and a single deletion of an LT gene often results in no cellular phenotype. To determine if the phenotype was unique to Δ*sleB*, we tested the effect of deleting the other predicted LT enzymes encoded in the genome, the membrane-bound MtlA and MltB (25). We readily generated the single *mltA* or *mltB* mutants, as well as the double deletion mutant, and assessed their growth profile. Interestingly, these strains did not exhibit impaired growth under regular growth conditions (Fig. S1). Thus, SleB appears to have more significant roles than the two Mlt enzymes under regular growth conditions.

### Δ*sleB* is sensitive to environmental stress

To further characterise the effect of deleting *sleB*, we monitored growth and cell division of Δ*sleB* cells by time lapse imaging. Overnight culture of *Z. mobilis* cells were inoculated on 1% agarose-pads made of the complex medium immediately before the imaging process started. While the wild-type ZM6 cells grew and divided without any defects on the agar-pad, Δ*sleB* cells exhibited a long lag-phase before the cells regrew (Fig. 3). During this lag phase, some cells lysed, as indicated by the change of their darkness by phase-contrast microscopy, but some Δ*sleB* cells eventually resumed growth (Fig. 3). These observations suggest that Δ*sleB* is sensitive to environmental changes, including the osmotic changes associated with placing the cells on agarose pads.

**Fig. 3.**
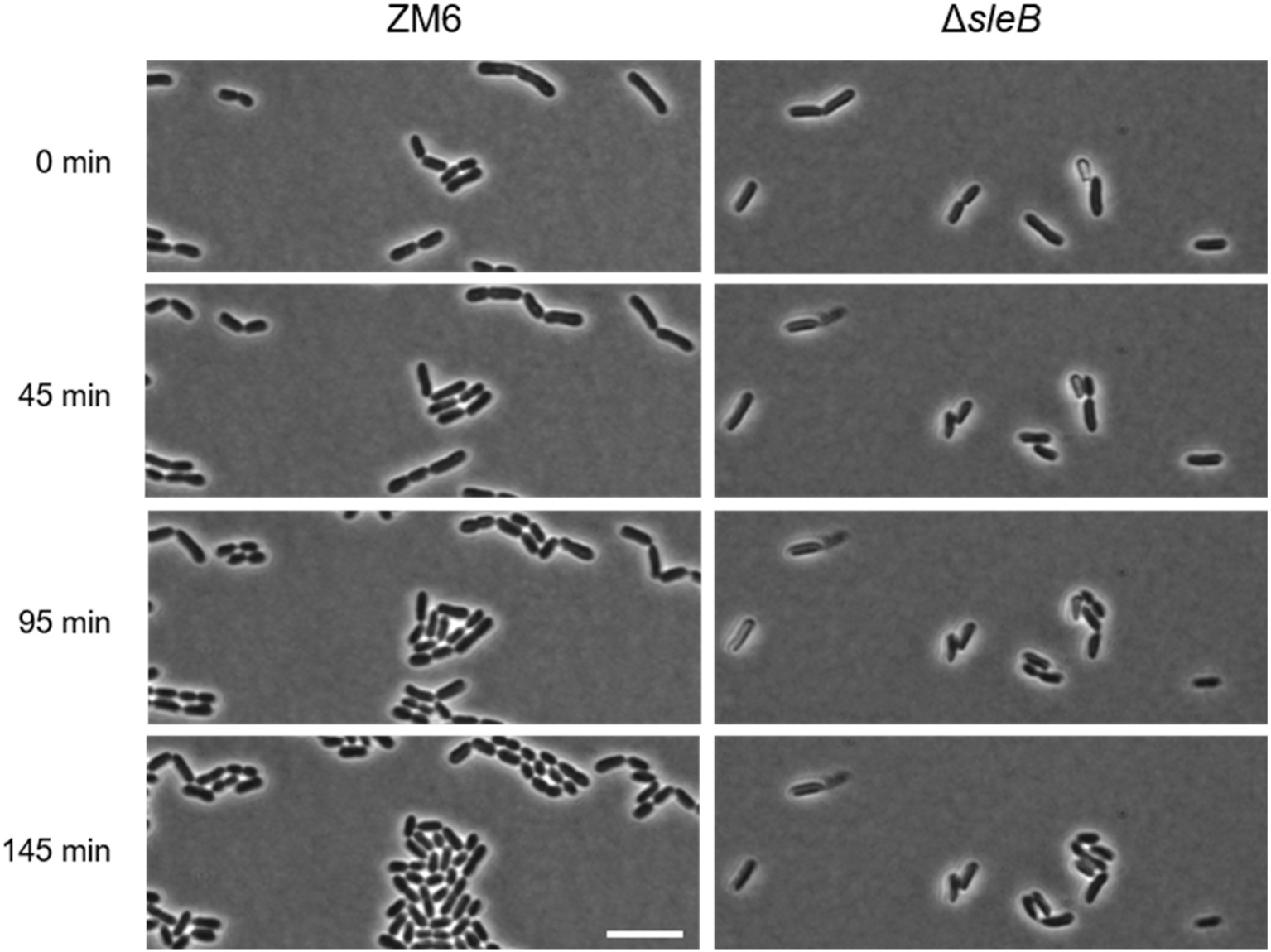
Time-lapse imaging of growing ZM6 and Δ*sleB* cells. A culture of fully grown *Z. mobilis* cells were spotted on the agarose-pad made of the regular complex medium before imaging started. Phase contrast images of ZM6 (left) and Δ*sleB* cells (right) were acquired at the time indicated in the figure. Scale bar, 10 μm.

### Δ*sleB* forms similar morphology as Δ*patA* under salt conditions

The growth profile and the time-lapse imaging experiments suggested that Δ*sleB* is sensitive to environmental stress, possibly through altered cell wall remodelling. This observation led us to examine whether Δ*sleB* is sensitive to salt. Because salt stress was shown to cause elongated cells with one wider cell pole (26), any malfunction in cell wall remodelling could produce a cellular phenotype under salt conditions. Also, Δ*patA* was previously shown to grow poorly and exhibit distinctive morphology under salt conditions, and thus, salt stress might be useful to assess potential functional link between SleB and PatA.

Interestingly, Δ*sleB* showed severe growth defects when stressed with salt. The final optical density of the Δ*sleB* culture was significantly lower than that of both the wild type ZM6 and Δ*patA* (Fig. 4A). Furthermore, the cell shape of Δ*sleB* was strikingly different from that of the wild type. While almost all of the wild-type cells exhibited a thick, bulged cell pole, Δ*sleB* cells were thinner and shorter (Fig. 4B). Interestingly, this cell shape resembles that of Δ*patA* cells under the same stress conditions (Fig. 4B) (17). Measuring the diameter of the Δ*sleB* cells showed that it was thinner than that of Δ*patA* cells, correlating with differences in the salt sensitivity (Fig. 4C). These results suggest that SleB might have a function related to PatA and PG O-acetylation.

**Fig. 4.**
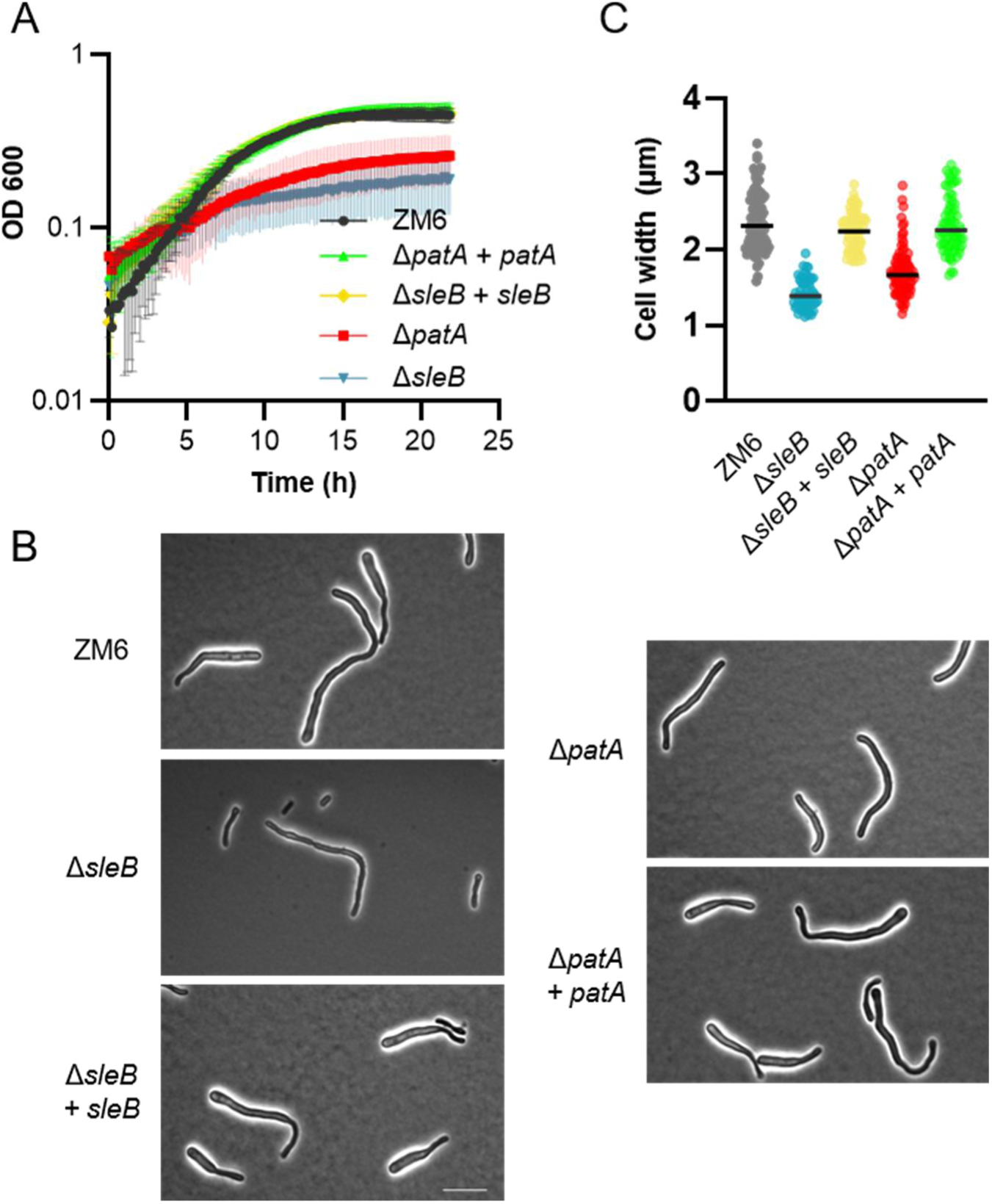
*Z. mobilis* Δ*sleB* cells exhibit impaired growth and morphology defects under salt (NaCl 225 mM) conditions. **(A)** Growth of *Z. mobilis* strains ZM6 (wild-type), Δ*sleB*, Δ*sleB* + *sleB, ΔpatA* and *ΔpatA + patA* under salt growth conditions. Biological replicates N = 3. The error bars represent the standard deviation. **(B)** Phase contrast images of ZM6, Δ*sleB* and Δ*sleB* + *sleB* cells grown for 14 hours under salt growth conditions. Scale bar, 10 μm. **(C)** A box-and-whisker plot presenting the maximum diameter of bulged cell pole of indicated *Z. mobilis* strains under salt conditions. Unpaired T-test was applied between strains, showing the significant difference between ZM6 and Δ*sleB* (P < 0.0001), Δ*patA* (P < 0.0001), Δ*sleB* + *sleB* (P = 0.007) and no significant difference between ZM6 and *ΔpatA + patA* (P = 0.3652). Biological replicates N = 3. Sample size, n = 87 cells (ZM6), n = 59 (Δ*sleB*) n = 74 (Δ*sleB* + *sleB*) n = 92 (Δ*patA*) n = 72 (Δ*patA* + *patA*).

### Δ*sleB* is sensitive to acetic acid stress

As Δ*sleB* appears to phenocopy Δ*patA*, we investigated other stresses, exogenous acetic acid and SDS, that produce a growth defect phenotype in Δ*patA* (Fig. 1).

Δ*sleB* did not grow at all in the presence of 0.0025% SDS, consistent with previous results showing that Δ*sleB* was sensitive against environmental stresses. Δ*sleB* grew poorer than wild-type under acetic acid stress in both liquid culture and on plate (Fig. 5). Notably, while Δ*sleB* displayed more severe growth defects than Δ*patA* under regular and stress (salt and SDS) conditions, both mutants exhibited a similar degree of sensitivity to acetic acid (Fig. 1 and 5).

**Fig. 5.**
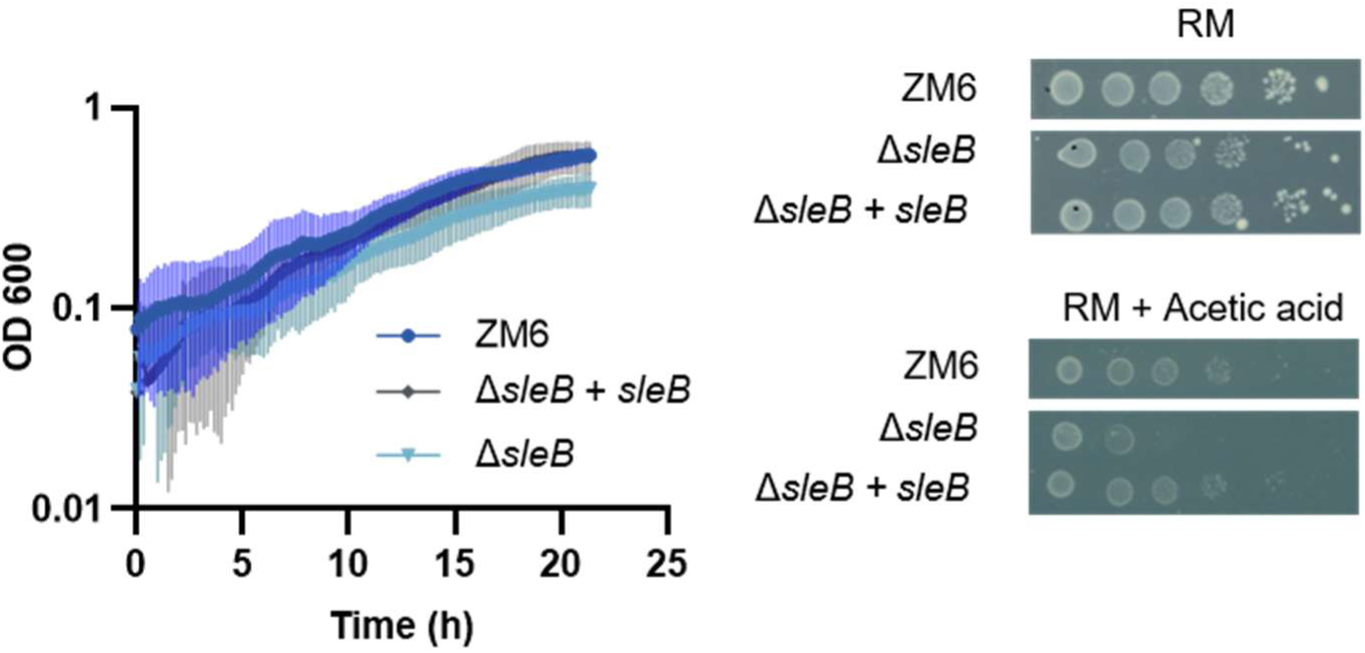
*Z. mobilis* Δ*sleB* cells show impaired growth under acetic acid (3.5 g/L) conditions. (Left) Growth of *Z. mobilis* strains ZM6 (wild-type), Δ*sleB* and Δ*sleB* + *sleB* under acetic acid growth conditions. Biological replicates N = 3. The error bars represent the standard deviation. (Right) Spot assay was performed to confirm the acetic sensitivity of Δ*sleB*. The image was taken after 96 hours. Biological replicates N = 3.

### Δ*sleB* cells have an altered PG composition

It was previously shown for *Neisseria meningitidis* that the truncation of the *ltgA* gene encoding an LT resulted in hyper O-acetylation of PG (27). The LT LgtA activates the PG O-acetyl esterase Ape, suggesting a mechanism in which de-O-acetylation by Ape facilitates subsequent cleavage of glycan chains by LgtA. Considering the gene proximity of *sleB* and the *pat* operon in the *Z. mobilis* genome, the lack of SleB might alter the O-acetylation level, resulting in the observed growth and stress phenotypes. Thus, we examined the PG composition of Δ*sleB*.

We purified PG from *Z. mobilis* cells and analysed the muropeptide composition as previously reported (Fig. 6A and Fig. S2) (17). The PG analysis method includes chromatography buffers adjusted to pH 6.0 that are sub-optimal for muropeptide separation but preserve the O-acetyl groups. Hence, chromatograms contain broader peaks compared to the standard *E. coli* profiles that do not account for O-acetylation (17, 28). The analysis revealed that Δ*sleB* had reduced TetraTetraAnh, which is an LT product (Fig. 6BC), and a non-significant reduction in the extent of O-acetylation (Fig. 6D). Interestingly, the proportion of dimeric muropeptides was significantly reduced from 40.1 % in wild-type to 29.0 % in Δ*sleB* (Fig. 6D). These changes in the muropeptide profile in the absence of SleB are likely relevant to the observed growth phenotypes.

**Fig. 6.**
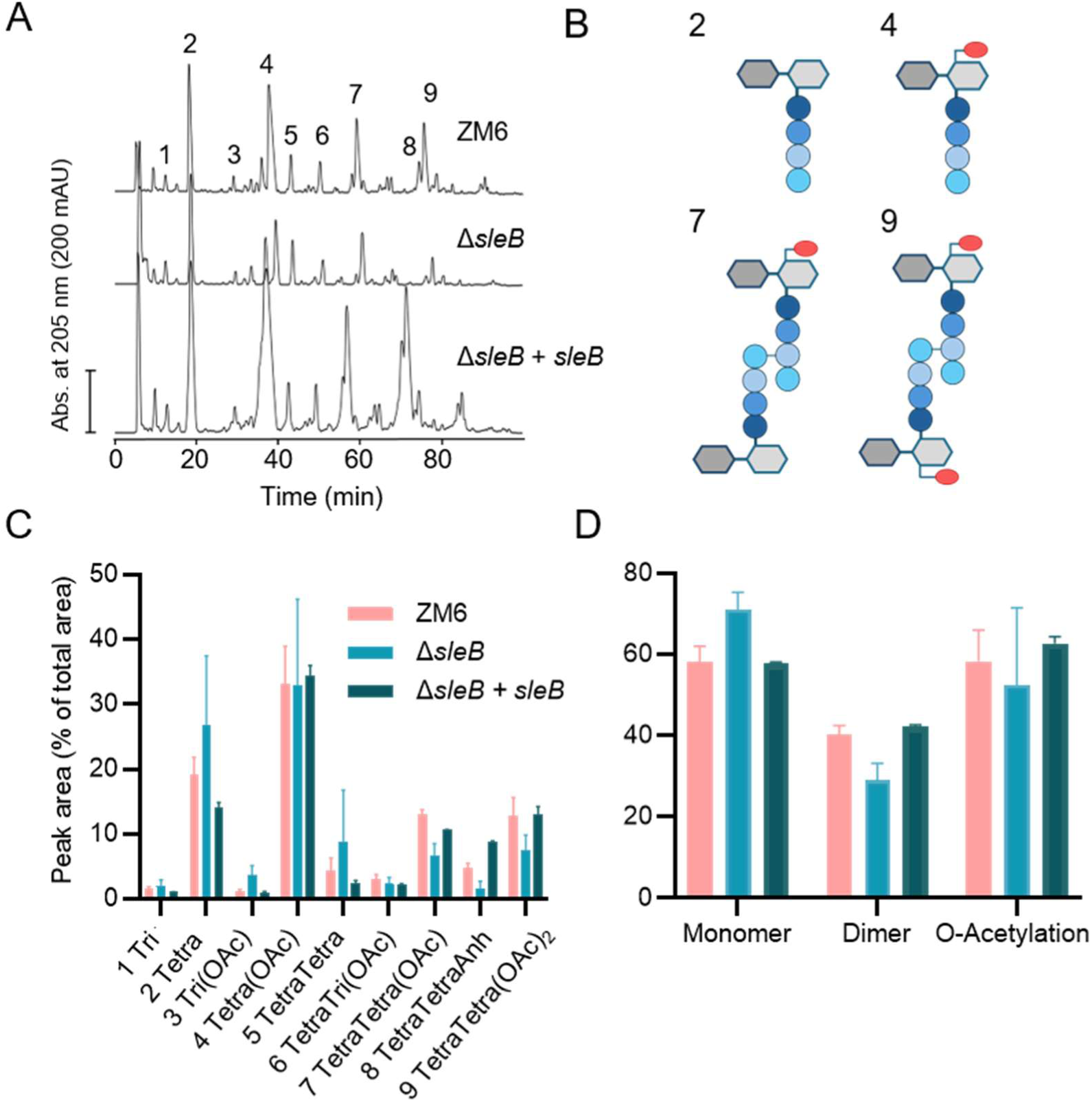
Muropeptide profile of ZM6 (wild type), Δ*sleB* and Δ*sleB* + *sleB* (complementation strain). **(A)** Cells of ZM6, Δ*sleB* and the *sleB* complementation strain were grown under regular growth conditions. The PG was isolated from growing cells and the muropeptides were released by cellosyl, reduced by sodium borohydride and separated by high-performance liquid chromatography (HPLC) under conditions that preserve O-acetyl groups. The indicated numbers represent identified individual muropeptides and their structures are shown in Fig. S2. **(B)** Structures of the major muropeptides released from *Z. mobilis PG*, illustrating the presence of O-acetyl groups at MurNAc (illustrated by red ovals) in some muropeptides. The structures are explained in Fig. S2. **(C)** Bar charts representing composition of individual muropeptides and **(D)** relative proportion of monomeric, dimeric and O-acetylated muropeptides from the measured strains. Biological replicates N = 2, for ZM6 and the complementation strain, N = 3 for Δ*sleB*. The error bars represent the variation.

### Overexpression of *sleB* also impairs growth

Next we aimed to determine the effect of *sleB* overexpression in *Z. mobilis*. First, we aimed to introduce an extra copy of *sleB* under its natural promoter, at a different locus using the homologous-recombination based method. However, we were not able to generate the strain, indicating that this would cause a loss of fitness or viability. We then used an alternative strategy, putting the *sleB* gene into the replicative plasmid pBBR under constitutively active *pdc* promoter (29). We were readily able to introduce the plasmid into the wild type ZM6 and Δ*patA*, and their growth was characterized under regular and salt conditions. The analysis showed that growth of both strain carrying pBBR *pdc-sleB* was not significantly perturbed under regular conditions (Fig. S3). However, under salt condition, both strains showed a higher sensitivity to the salt than the control strain that carry the empty pBBR plasmid (Fig. S3).

## Discussion

SleB was previously characterised as a **<u>s</u>**pore cortex **l**ytic **<u>e</u>**nzyme (Sle) important for germination of spores in spore-forming, Gram-positive bacteria (30–32). The crystal structure of SleB from *Bacillus cereus* shows a catalytic domain with a similar fold to that of the well-studied Slt70 from *Escherichia coli*, despite limited amino acid conservation between both proteins (33). The *sleB* gene appears to be conserved across alpha-proteobacterial species (5), but its biological role has remained unexplored.

*Z. mobilis* Δ*sleB* exhibited impaired growth even under favourable growth conditions and severe sensitivity to environmental stress consistent with cell envelope defects. The pronounced phenotypes resulting from the deletion of a single LT gene are rather unique, because, in *E. coli*, triple LT gene deletions of *mltA*, *mltB*, and *slt70* did not lead to detectable growth defects (25) while six LT-encoding genes were shown to be dispensable for normal cell growth in *Vibrio cholera* (6, 34). Similarly, the triple mutant of soluble LTs, Δ*sdpABC* exhibited only minor growth and morphological defects in *Caulobacter crescentus* (35). The significant impact of *sleB* deletion may be due to the small size of the *Z. mobilis* genome arrangement which appears to have less functional redundancy (36).

The *Z. mobilis sleB* mutant cells displayed thin and long morphology, with occasional formation of cell chains with thread-like structures between the two daughter cells under regular growth conditions (Fig. 2). This is consistent with previous studies showing that some LTs function in daughter cell separation in Gram-negative bacteria (3). *Z. mobilis* SleB might localise at the septum and cleave the PG during cell division. We attempted to determine the sub-cellular localization of SleB, however, N- or C-terminal sfGFP fusions were not functional.

In *Pseudomonas aeruginosa*, the LT RlpA has been shown to contribute to daughter cell separation by degrading denuded glycan strands, which are generated by amidase activity at the septum (37). RlpA contains a SPOR domain responsible for binding to denuded peptidoglycan and promoting localisation of RlpA at the division site (38). In contrast, soluble SleB lacks a SPOR domain and might recognise its substrate by a different mechanism. A previous study showed that *Bacillus anthracis* SleB binds to a spore specific modification, muramic-δ-lactam, to recognise and degrade spore cortex PG for germination (30). Thus, further studies need to identify the PG-binding mechanism of *Z. mobilis* SleB, assess the enzymatic activity of SleB on substrate PG with or without peptide stems, and determine protein interaction partners of SleB.

The muropeptide profile of Δ*sleB* PG possesses a significantly reduced proportion of cross-linked muropeptides comparing to the wild-type under regular growth conditions. This ‘relaxed’ PG might be an underlying cause for the weakened cell envelope of Δ*sleB*. The observed reduction of cross-linkage is intriguing, as we previously overexpressed the endopeptidase MepM in *Z. mobilis* and observed no significant reduction in the dimer proportion (17).

Interestingly, Δ*sleB* exhibited similar morphology to Δ*patA* under salt conditions. Salt stress severely disturb cell division of *Z. mobilis* by unknown mechanism, causing the cells to elongate and develop one wider, bulged pole (17). The thin morphology indicate that SleB is involved in additional PG related functions beyond daughter-cell separation. Furthermore, both mutants showed comparable sensitivity to acetic acid. Importantly, under the tested condition acetic acid is not dissociated and thus can diffuse across the cell membrane, and subsequently dissociate in the cytoplasm where the pH is neutral. Furthermore, acetic acid stress was the only condition under which cell growth of Δ*patA* and ΔsleB were comparable. These observations indicate that the two mutants might share a physiological disturbance caused by intracellular acidification or other cytoplasmic malfunction caused by external acetic acid.

The LT LtgA in *Neisseria meningitidis* was previously shown to contribute to modulation of PG O-acetylation (27). Strikingly, truncation of LtgA resulted in the hyper O-acetylation of PG and defects in cell morphology. The underlying mechanism appears to involve compromised activation of the PG esterase Ape1 which removes O-acetyl modifications from the cell wall (27). Considering the gene landscape of *sleB* that is adjacent to *pat* operon which includes *ape* in *Z. mobilis*, it is tempting to speculate that SleB might be involved in maintenance of basal PG O- acetylation level in a similar manner to LtgA. Our muropeptide analysis revealed only minor, if any, differences of O-acetylation between the wild-type and Δ*sleB*. However, the analysis represents a ‘snapshot’ of the overall PG composition, and the potential involvement of SleB in O-acetylation may not be ruled out. One possibility is that the overall rates of PG O-acetylation and de-O-acetylation might be altered in Δ*sleB*. Previously, the depletion of *ape* by CRISPRi caused significantly impaired cellular fitness in *Z. mobilis* type strain ZM4 (36), and our attempts to delete *ape* in the strain ZM6 were unsuccessful. These observations suggest that hyper O-acetylation of PG, caused by the lack of Ape activity, may be toxic in *Z. mobilis*. This contrasts with other species in which *ape* was dispensable and the elevated O-acetylation level did not cause severe growth defects (27, 39). Thus, *Z. mobilis* might possess an unidentified mechanism that prevents excessive PG O-acetylation by modulating the activity of the O-acetylation machinery, and the impact of LT deletion on PG O- acetylation may differ from that in other bacteria with PG O-acetylation. Alternatively, O-acetylation may be spatially regulated within the PG and this pattern might be disrupted in the absence of SleB. More work is needed to gain a better understanding of the dynamics and spatial control of O-acetylation in *Z. mobilis* and other bacteria.

## Acknowledgements

This work was supported by Newton International Fellowship (The Royal Society) to K.F. (NIF\R1\221190).

## Notes

### Competing Interest Statement

The authors have declared no competing interest.

