## Supplemental Figure S1-S3 and Supplemental table S1 -2 for "Hypothetical lytic transglycosylase SleB is important for cell fitness in *Zymomonas mobilis*"

**Supplemental Figures**

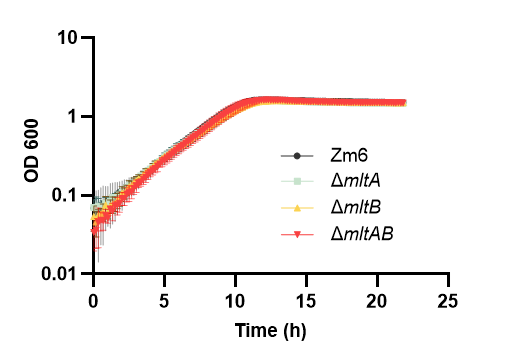

**Fig. S1.** Growth of *Z. mobilis* strains ZM6 (wild-type), Δ*mltA*, Δ*mltB* and Δ*mltAB* under regular growth conditions. Biological replicates N = 3. The error bars represent the standard deviation.

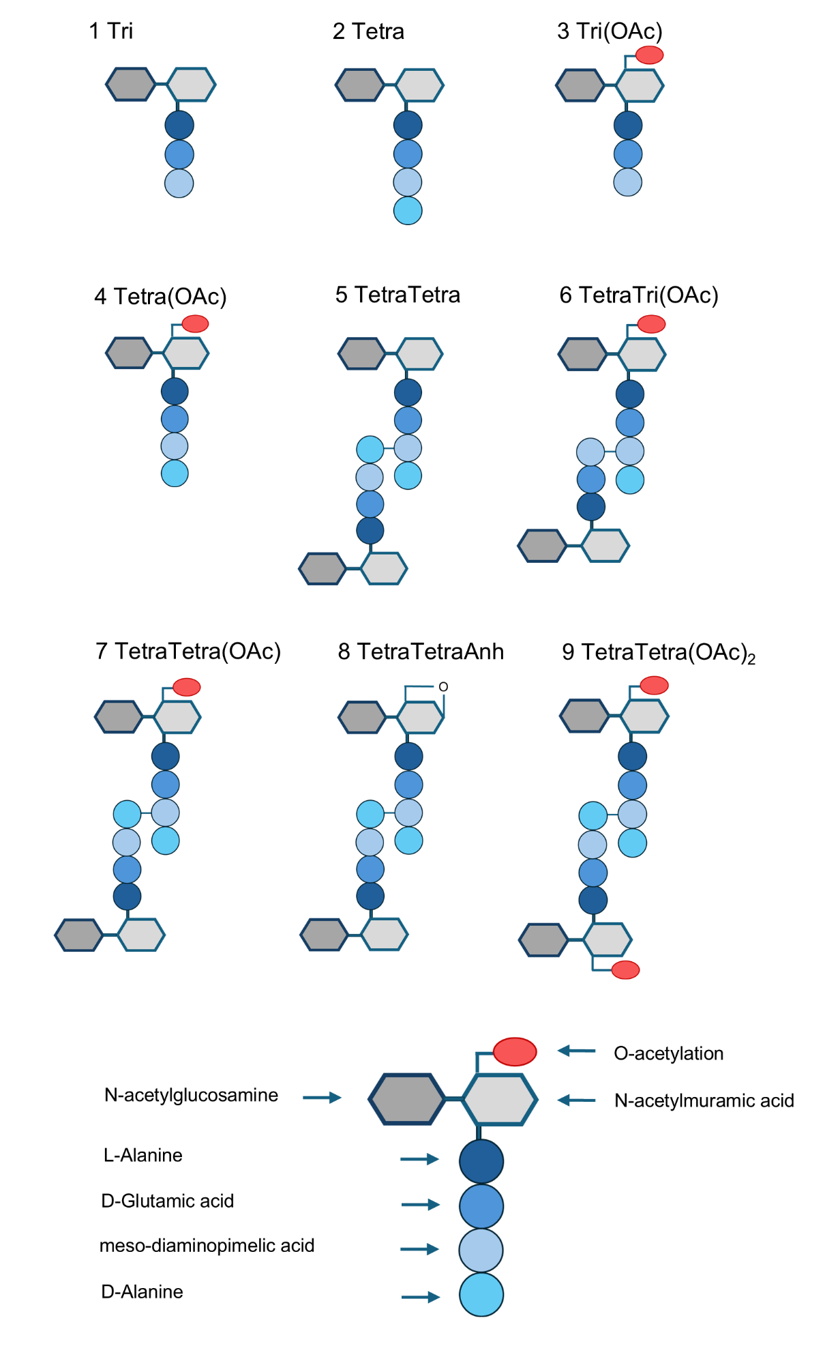

**Fig. S2.** Schematic structures of muropeptides released from *Z. mobilis* peptidoglycan. The number corresponds to peak numbers in Fig. 6A.

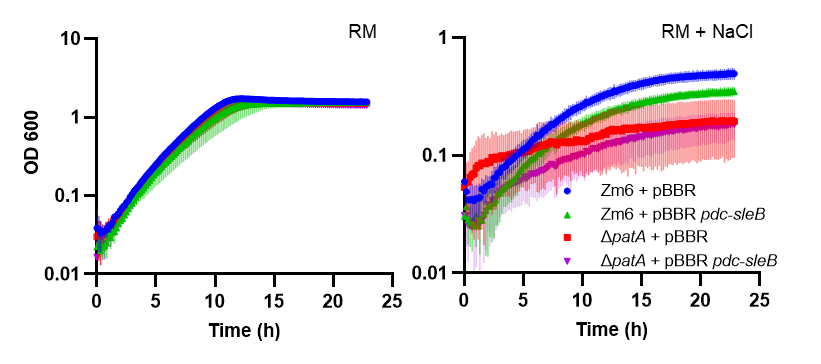

**Fig. S3**. Overexpression of *sleB* impairs growth in *Z. mobilis*. (Left) Growth of *Z. mobilis* strains ZM6 + pBBR, ZM6 + pBBR *pdc-sleB*, Δ*patA* + pBBR, Δ*patA* + pBBR *pdc-sleB* under regular growth conditions. (Right) Growth of the same strains under salt growth conditions (NaCl 225 mM). For the plasmid maintenance, kanamycin (200 μg/mL) was added in the growth medium.

**Supplementary tables**

**Table S1.** Bacterial strains and plasmids used in this study.

| Bacterial strains | |  |  | Genotype/ description/reference |
| --- | --- | --- | --- | --- |
| *Zymomonas mobilis* ZM6 | | |  | ATCC29191 purchased from DSMZ |
| *Escherichia coli* Dh5a | |  |  | Cloning strain (Lab stock) |
| *Escherichia coli* WM6026 | | |  | Conjugation strain (1) |
| *Zymomonas mobilis* + SacB | | |  | *B. subtilis* sacB at ZZ*6*_*1449* (2) |
| *Zymomonas mobilis* Δ*patA* | | |  | Δ*ZZ6_0812* (2) |
| *Zymomonas mobilis* Δ*patA + patA* | | | | Δ*ZZ6_0812* + *ZZ6_0812* (2) |
| *Zymomonas mobilis ΔsleB* | | |  | *ΔZZ6_0810* (2) |
| *Zymomonas mobilis ΔsleB + sleB* | | |  | *ΔZZ6_0810 + ZZ6_0810* (this study) |
| *Zymomonas mobilis ΔmltA* | | |  | *ΔZZ6_1479* (2) |
| *Zymomonas mobilis ΔmltB* | | |  | *ΔZZ6_0234* (2) |
| *Zymomonas mobilis ΔmltAΔmltB* | | |  | *ΔZZ6_0234 - ΔZZ6_1479* (this study) |
| *Zymomonas mobilis* pBBR *pdc-sleB* | | | | pBBR + *pdc-sleB* (this study) |
| Plasmids |  |  |  | description/reference |
| pPK15534 |  |  |  | Suicide vector (1) |
| pPK15534 *+ sacB* | |  |  | pPK15534 carrying *B. subtilis sacB* (this study) |
| pPK15534 *+ sacB ΔZZ6_0810* | | |  | pPK15534 + *sacB* carrying Δ*ZZ6_0810* cassettes (2) |
| pPK15534 *+ sacB + ZZ6_0810* | | |  | pPK15534 + *sacB* carrying *ZZ6_0810* (this study) |
| pPK15534 *+ sacB ΔZZ6_1479* | | |  | pPK15534 + *sacB* carrying ΔZ*Z6_1479* cassettes (2) |
| pPK15534 *+ sacB ΔZZ6_0234* | | |  | pPK15534 + *sacB* carrying Δ*ZZ6_0234* cassettes (2) |
| pBBR *+ pdc-sleB* | |  |  | pBBR carrying *pdc-sleB* (this study) |

**Table S2.** Oligonucleotides used in this study.

| Oligo | Sequence | | Use |
| --- | --- | --- | --- |
| **NKF99** | acccgtggttcatgcatcagcgtatggggctgacttcaggtgc | | ***sleB complementation*** |
| **NKF100** | acgccttttctagcaaaggggtaattctcatgtttgacagcttatcac | |  |
| **NKF219** | cacctgaagtcagccccatacgctgatgcatgaaccacgggtg | |  |
| **NKF224** | gctgtcaaacatgagaattacccctttgctagaaaaggcgtgcc | |  |
| **NKF267** | aatggcaccgacctttgatagcgtcgttttagttatatcttggcttgctc | |  |
| **NKF268** | gccaagatataactaaaacgacgctatcaaaggtcggtgccattatc | |  |
| **NKF269** | tagggggtataatccggtctcaacatcgctattggaagaagctggaag | |  |
| **NKF270** | agcttcttccaatagcgatgttgagaccggattataccccctaggaac |  |  |
| **NKF340** | aatcatgtctagattcaaggtgtcccgttcctttttccc | | ***pdc*-*sleB*** |
| **NKF341** | tcctaaagcggacattgcttactccatatattcaaaacactatgtctg | |  |
| **NKF342** | tatatggagtaagcaatgtccgctttaggacatttatttcgtaaaaaatg | |  |
| **NKF343** | aatcatgctcgagttaacgatagaaaacgtggctacccaaagc | |  |
